# The indispensable soma of *Cardiocondyla obscurior* ants

**DOI:** 10.1101/2022.10.02.510526

**Authors:** LM Jaimes-Nino, A Süß, J Heinze, E Schultner, J Oettler

**Affiliations:** Zoologie/Evolutionsbiologie, Universität Regensburg, 93053 Germany

**Keywords:** Aging, Continuusparity, Social insects, Worker lifespan

## Abstract

The evolutionary mechanisms that shape aging in social insects are not well understood. It is commonly assumed that queens live long and prosperous, while workers are regarded as a short-lived disposable caste because of their low reproductive potential. Queens of the ant *Cardiocondyla obscurior* gain high fitness late in life by increasing investment into sexual offspring as they age. This results in strong selection against senescence until shortly before death. Here, we show that workers have the same lifespan and shape of aging as queens, even though workers lack reproductive organs and cannot gain direct fitness. Under consideration of the prevailing aging theories and the biology of the species, we hypothesize that programmed aging has possibly evolved under kin selection.

**Impact statement:** Morphologically distinct fertile queen and sterile worker castes in the model ant *Cardiocondyla obscurior* show the same pace and shape of aging, contradicting the paradigm of queen/worker lifespan divergence in social insects.

## Introduction

Queens of some social Hymenoptera (ants and bees) live long while being highly fertile, seemingly avoiding a trade-off between lifespan and reproduction (Hartmann and Heinze, 2003; Korb, 2016; Korb et al., 2020; Parker, 2010). Using the tiny ant *Cardiocondyla obscurior,* a model for social insect aging (reviewed in Oettler and Schrempf, 2016), we recently identified molecular processes associated with queen aging (Wyschetzki et al., 2015), and demonstrated that the strength of selection on age-biased genes differs between social and solitary insects (Harrison et al., 2021). Based on these insights and lifespan data from *C. obscurior* queens, including transcriptomic data from queens shortly before death, a life history framework was proposed (Jaimes-Nino et al., 2022).

The concept of “continuusparity” is grounded in a pattern observed in many ants, namely that resources are first invested into workers, and only when colony size has reached a certain threshold, resource investment is diverted to the production of sexual offspring (males, queens) (Oster and Wilson, 1978). Thus the strength of selection against senescence does not decline with age despite life-long reproduction (Jaimes-Nino et al., 2022). Irreversible reproductive division of labor between queens and workers is a pre-requisite for the evolution of continuusparity, however, not all social insects must exhibit continuusparity, for example, species with only one caste. In respect to senescence, continuusparous social insects should sit between iteroparous and semelparous species, the former experiencing decreasing selection strength after the first reproductive bout, while the latter undergo strong selection against senescence until a single, and final, reproductive bout, followed by reproductive death.

As progress is made in understanding aging in ant queens, the aging patterns of workers remain largely unknown. Anecdotal references such as “mother queens live much longer than workers in all groups of ants” (Hölldobler and Wilson, 1990), and “all eusocial taxa show a divergence of long queen and shorter worker life spans, despite their shared genomes and even under risk-free laboratory environments” (Kramer et al., 2022) have been made. However, studies of worker lifespan lacked age-controlled cohorts (Chapuisat and Keller, 2002; Gordon and Hölldobler, 1987; Modlmeier et al., 2013; Negroni et al., 2021; Schmid-Hempel and Schmid-Hempel, 1984), surveyed marked individuals in the field without distinguishing between extrinsic and intrinsic mortality (Calabi and Porter, 1989; Gordon and Hölldobler, 1987; Schmid-Hempel and Schmid-Hempel, 1984), or monitored lifespan of temperate species with artificial hibernation (Kramer et al., 2016). Furthermore, it is difficult to find reliable lifespan data of queens for comparison (Kramer and Schaible, 2013a). Here, we present the first empirical study of worker lifespan under controlled conditions in an ant with morphologically distinct castes and show that the paradigm of lifespan divergence between ant castes is not true.

## Methods

*Cardiocondyla obscurior* (Formicidae: Myrmicinae), the two-colored heart node ant, is a small tramp species (queens: 3mm, workers: 2mm, (Seifert, 2003), which forms colonies comprising a few queens and several dozen workers in trees and shrubs around the tropics and subtropics (Heinze, 2017; Oettler, 2021). Queens and workers differ in discrete traits such as the presence of wings and ocelli, and, importantly, workers in this species lack reproductive organs and are thus fully sterile. Queens produce on average 300 (± 198 SD) offspring over 25 (± 8 SD) weeks (Jaimes-Nino et al., 2022) and have control over caste and sex allocation (De Menten et al., 2005; Schultner et al., 2021).

We obtained lifespan data for worker ants by setting up colonies containing 10 dark-colored worker pupae (a few days before hatching into adults) from independent stock colonies of a laboratory population collected in 2011 in Japan ("OypB", (Errbii et al., 2021; Schrader et al., 2014; Ün et al., 2021). Two virgin queens were added to the colonies to help the workers eclose and were removed after two weeks. Colonies were kept in climate-controlled conditions (26°C/22°C, 12h/12h light, 75-80% humidity) in nests made of acrylic glass with narrow slits, sandwiched between two microscope slides and covered with a dark foil, placed in a square petri dish half-filled with plaster. The ants always had access to water, and were fed with honey, *Drosophila* and pieces of cockroaches twice per week. To test for an effect of workload on worker lifespan, colonies were subjected to one of three treatments: 1) no larvae (NL), 2) with two second instar larvae (low workload, LW), and 3) with ten second instar larvae (high workload, HW) (n=20 each). To keep workload constant, in the two treatments containing brood, colonies were checked every few days and larvae that had pupated were removed and replaced by new second instar larvae.

After 18 weeks there was no difference in survival between workers with or without larvae (see results). This led us to adjust treatments to provoke more variation in lifespan. Colonies were standardized to five workers, and those containing fewer than five workers were excluded (n after exclusion: NL=18, LW=20, HW=16). The remaining replicates from each treatment were split into two groups; in one group, workers were kept without larvae while in the other, two second instar larvae were added to simulate low workload (n after split: NL=9, NL-LW=9, LW-NL=10, LW=10, HW-NL=8, HW-LW=8). Larvae were replaced as described above. As after 36 weeks still no effect of workload treatment was apparent (see results), we removed all larvae and continued to monitor worker survival weekly until all workers had died.

Differences in survival across worker treatments and between workers and queens (using queen data from Jaimes-Nino et al., 2022a) were tested. A Cox proportional hazard mixed-effect model was implemented (coxme package in R, v. 2.2.-16) using colony as a random factor; post-hoc tests using multiple comparison of means were run where appropriate (Tukey contrasts, *glht* function in multcomp package in R, v. 1.4-19). Relative mortality as a function of age was mean-standardized by dividing age-specific mortality by its mean (following Jones et al. 2014). We considered mortality trajectories from the age at eclosion to the age at which 5% survivorship from eclosion occurs. Predictions of the data were visualized using the loess method with the geom_smooth function and default span (ggplot2 v.3.3.2). The mean life expectancy of all workers was calculated with the *survfit* function (survival package in R, v.3.3-1) using Kaplan Meier survival analysis.

## Results

After 18 weeks, there was no difference in worker survival between the treatments with and without larvae (Fig 1, coxme, NL vs. LW, z-value=−1.35, p=0.37, NL vs. HW, z-value=1.35, p=0.37), but survival of workers with low workload was higher than survival of workers with high workload (coxme, LW vs. HW, z-value=2.7, p=0.02). Shortly after colonies were standardized to five workers and split into groups without larvae or with low workload (week 18), survival differed between treatments (coxme anova, *X*^2^_5_=14.24, p<0.05), but this difference was not significant after correction for multiple testing. After 36 weeks there was still no effect of the presence of larvae on worker survival (coxme, NL vs. NL-LW, z-value=−1.08, p=0.89, NL vs. LW, z-value=−0.79, p=0.97, NL vs. HW-LW, z-value=1.58, p=0.61), so we stopped replacing larvae, removed any remaining brood items and continued to monitor survival weekly. As some larvae were potentially overlooked and may have developed into adult workers, 14 replicates (NL-LW=8, LW=3, HW-LW=3) were discarded at this point. The remaining replicates were monitored until the last worker was recorded alive after 50 weeks (Fig 2A).

**Figure 1.**
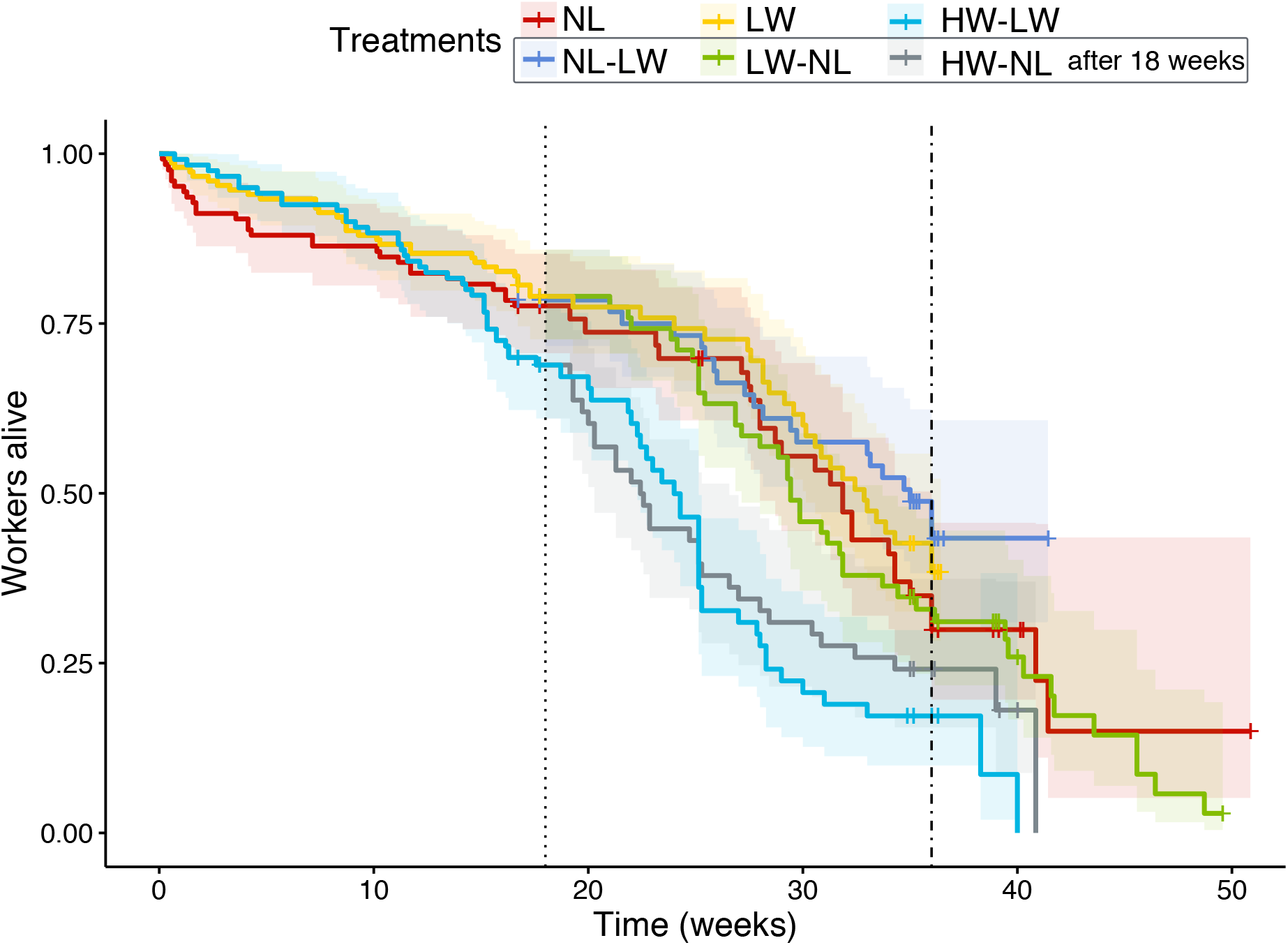
Survival probability curve of *C. obscurior* workers depending on treatment (NL: No larvae; LW: Low workload; HW: High workload) estimated using Kepler-Meier analysis. At week 18, indicated by the dotted line, colonies were standardized to 5 individuals and half of the colonies from the NL and LW treatments, and all HW colonies, were subjected to a treatment change (from NL to LW: NL-LW, from LW to NL: LW-NL, from HW to NL and LW: HW-NL, HW-LW). After 36 weeks, indicated by the dot-dashed line, no more larvae were added. The NL-LW and LW curves are truncated after 36 and 42 weeks due to the dismissal of replicates in which larvae may have eclosed.

**Figure 2.**
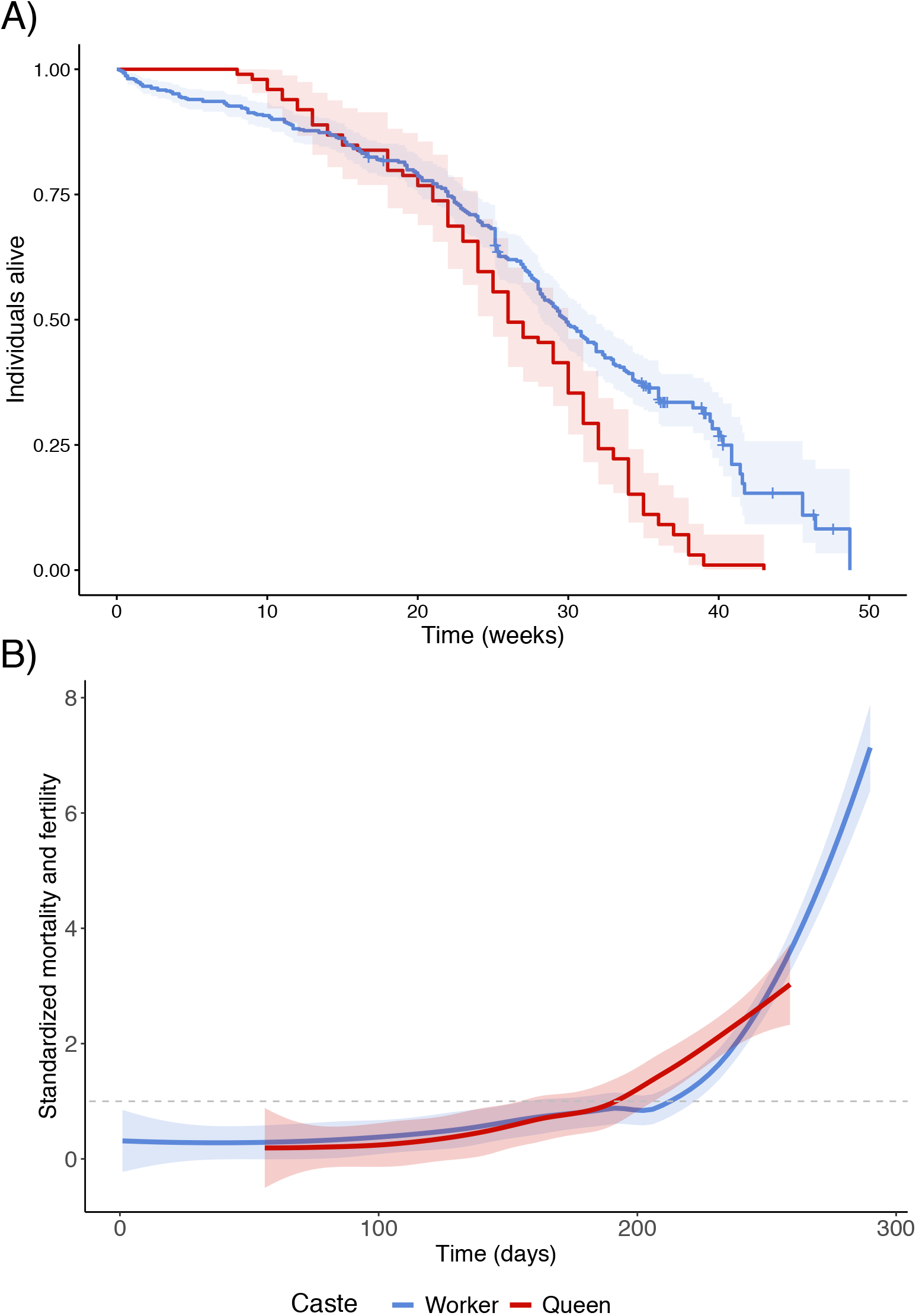
Worker and queen survival and standardized mortality. A) Survival probability curve of queens (n = 99; from Jaimes-Nino et al. 2022) and workers (n=530). B) Relative mortality as a function of age. Mortality is standardized by dividing age-specific mortality of queens and workers by their means after eclosion (following Jones et al., 2014). The graph uses a Loess smoothing method (span = 0.7) and a confidence interval of 95%. The dashed gray line at y = 1 indicates when relative mortality is equivalent to mean mortality.

Median worker lifespan over all treatments was 29 weeks (209 days, CI 95%=198-223 days).

The lifespans of workers were similar to those of queens obtained in a recent study using similar laboratory conditions (Jaimes-Nino et al., 2022). Workers had a lower hazard rate compared to queens (Hazard ratio = 0.55, coxme anova, *X*^2^_1_=56.95, p<0.001). Strikingly, the shape of aging, i.e. how mortality increases with time, was similar in the two castes, with mortality below average before day 200, and thereafter continuously increasing (Fig 2B).

## Discussion

### No causal relationship between fertility and lifespan

Aging is linked to metabolic rate, growth, and reproduction, and determined by life-history optimization (White et al. 2022). The shape of *C. obscurior* queen aging is characterized by an increasing investment into sexual reproduction with age, and thus a higher probability for (sexual) offspring to come into existence (Hamilton, 1966). After this bout of sexual production, mortality increases. Queens die soon after they cease egg laying, suggesting that reproduction is optimized and intrinsic resources are depleted (Jaimes-Nino et al., 2022). Finding a similar pace and shape of aging in workers is surprising because queens and workers differ in morphology (Fig 3, (Oettler et al., 2019), holobiome (Klein et al., 2016), presumably extrinsic mortality, and, perhaps most importantly, reproductive physiology and behavior and their associated energy expenditures.

**Figure 3.**
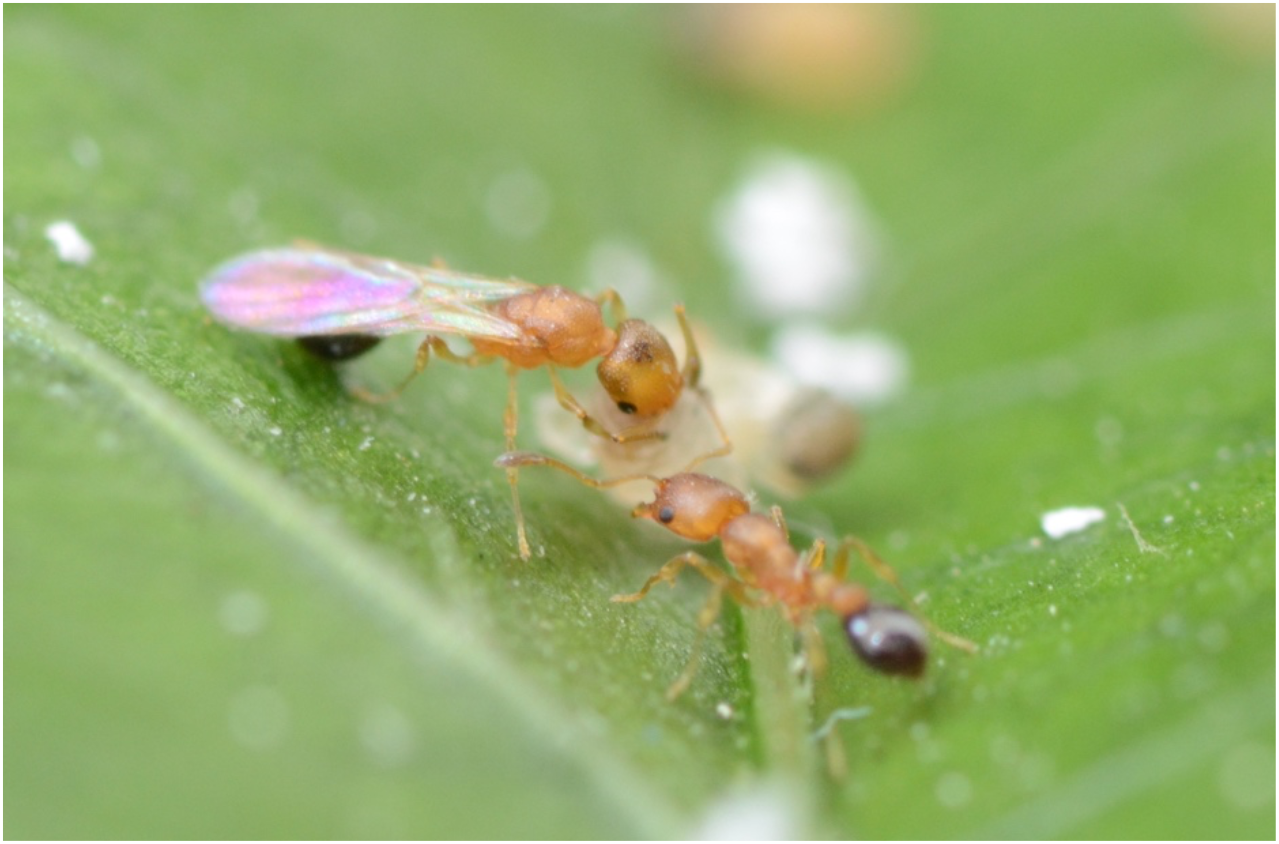
A virgin *C. obscurior* queen (left) and a worker tending to some brood. © Lukas Schrader.

Fertility is the most common proximate explanation for long lifespans in social Hymenoptera. Several studies have suggested causality between lifespan variation and core gene pathways and proteins involved in reproduction such as TOR, insulin-like signaling, juvenile hormone, and vitellogenines (honeybees: (Corona et al., 2007; Münch and Amdam, 2010; Rueppell et al., 2016; Seehuus et al., 2006); ants: (Korb et al., 2020; Negroni et al., 2021; Yan et al., 2022). In *C. obscurior*, queen fertility is positively correlated with lifespan (Heinze and Schrempf, 2012; Jaimes-Nino et al., 2022; Kramer et al., 2015), but the two traits are not causally linked: mated queens and queens mated to sterilized males lived longer than unmated queens, although the latter two both had low fertility and only produced unfertilized, haploid male eggs (Schrempf et al., 2005). Mated and sham-mated queens also differed from unmated queens in cuticular hydrocarbon composition and gene expression (Will et al., 2012; Wyschetzki et al., 2015). Together, this points toward a causal relationship between aging rate and mating status, rather than fertility. Fertility and fecundity are certainly not linked to the lifespan of workers because *Cardiocondyla* workers have no ovaries. This demonstrates that reproduction and lifespan are regulated independently in this ant, and supports a growing number of proximate and pathway-oriented studies in solitary animals showing that these two fundamental life history traits can be uncoupled (Dillin et al., 2002; Lind et al., 2021; Mason et al., 2018).

### Queen and worker lifespans are similarly long and variable

Worker lifespan is probably as variable as queen lifespan, which in addition to mating status responds to social environment (Schrempf et al., 2011, 2005), mating partner (Schrempf and Heinze, 2008) and immunity (Schrempf et al., 2015; Ün et al., 2021). The lifespans of queens reported in Jaimes-Nino et al. (2022) exceeded those in previous studies likely because laboratory conditions changed after 2015 (nest architecture, temperature, humidity, feeding rate). In the present study, workers also lived considerably longer than in the previous experiment in which focal workers were kept with 10 or 20 workers (each marked by tarsal clipping), and a fertile queen, presumably making life more demanding (Jaimes-Nino et al., 2022). In this study, removal of larvae after 18 weeks led to a temporary drop in the survival curves. This could point to a social or physiological factor that affects worker behavior, leading to more search activity (“where is the brood?”), and resulting in premature death (Koto et al., 2015). The slightly lower hazard rate in workers compared to queens may be explained by different energy requirements at the end of life.

The presumed universality of queen/worker lifespan divergence in ants, if true (~130MYR evolution, ~14.000 species), provokes the question why queens do not live longer (or workers shorter) in *C. obscurior*. While it seems futile to speculate without more data from more species for perspective, one explanation may be that this ant does not display traits thought to be associated with the evolution of lifespan divergence between castes, including extreme size polyphenism, colonies headed by single, highly fertile queens, and large colony sizes (Kramer et al., 2022; Kramer and Schaible, 2013b). It has also been argued that in species with multiple queen colonies such as *C. obscurior*, new cohorts of workers produced by young queens can outnumber older workers, decreasing the average relatedness of workers to the previous generations of queens. This increases the risk of queens being “dismissed” earlier by younger worker cohorts and may select for shorter lifespans in queens of polygynous species (Boomsma et al., 2014; Kramer et al., 2022). However, *C. obscurior* shows no signs of conflict between castes over sex or caste allocation (De Menten et al., 2005; Schultner et al., 2021) in contrast to many other ants (Heinze, 2004), speaking against this adaptationist explanation.

## Conclusions

Regarding the different approaches to gerontology, what can we learn from this ant? A first approach focuses on aging from a damage-based perspective and asks why evolution, which managed to make life in the first place, is not capable of maintaining life for eternity (minus some entropy). Research is usually directed at mechanisms underlying lifespan variation within species, and indeed there is overlap across taxa (e.g. dietary restriction, Fontana et al., 2010). *C. obscurior,* with its morphologically and physiologically distinct queen and worker castes, which nevertheless display a similar pace and shape of aging, is a unique model in this context (Jones et al., 2014), especially because proximate aging patterns contrast partly with those found in fruit flies (Harrison et al., 2021; Wyschetzki et al., 2015). While a senescent phase is only brief, this diphenic model will allow for disentangling how resources and energy are allocated into growth, aging and reproduction.

A second approach adds an evolutionary perspective, and considers life history strategies and putative associated metabolic or functional trade-offs (i.e. between lifespan and reproduction) to underlie senescence. Empirical evidence for this is found with mixed success (Dillin et al., 2002; Lind et al., 2021; Maklakov and Chapman, 2019; Mason et al., 2018). The results presented here do not support the idea that fertility-related trade-offs affect worker lifespan, simply because workers do not reproduce. There is also no evidence of fertility/lifespan trade-offs in queens (Jaimes-Nino et al., 2022; Schrempf et al., 2017), thus theories implying trade-offs do not apply here, including e.g., antagonistic pleiotropy, which postulates that genes with positive effects on the germline in early life can be selected for even if they cause senescence in the soma later in life. As antagonistic pleiotropy requires a phase after the point of strongest selection, it is unlikely to be effective in *C. obscurior* because there is no such phase (Jaimes-Nino et al., 2022). Along the same lines, metaphorically speaking, the results suggest that *C. obscurior* workers are not a disposable soma.

In contrast to the first two approaches aimed at understanding senescence, a third controversial perspective considers whether death is not just a consequence of lifelong wear and tear, and lifespan a trait under selection. The idea of programmed aging is strongly contested for insoluble problems with its logic (Kirkwood and Melov, 2011; Kowald and Kirkwood, 2016). First and foremost, how could selection ever favour death over reproduction, without invoking group selection? And why should aging be programmed in the first place, to shorten generation time for the betterment of the species? Programmed aging is also difficult to envision in iteroparous species exhibiting repeated reproduction and increasing risks and costs of somatic maintenance, and a senescent phase that is rarely expressed in nature. This is partly a problem of semantics because “iteroparity” comprises a wide range of reproductive strategies, including humans with an exceptionally long post-reproductive and senescent phase. The concept of programmed aging seems easier to imagine in semelparous species, where reproduction is a single event. However, in semelparous species it is not lifespan that is determined. Instead, reproduction is optimized towards one episode, which is often triggered by season, probability of encountering prey or hosts, or other extrinsic factors.

Despite all valid counterarguments, including inconceivable proximate mechanisms, programmed aging can explain not only the shape of aging of *C. obscurior* queens, but also the similarity of queen and worker aging patterns. All queens show increasing investment into sexuals with age, irrespective of overall fertility, lifespan or colony size, followed by increasing mortality. This indicates some sort of Zeitgeber, informing the queen to increase investment into sexual production. As potential environmental signals are absent under controlled conditions in the lab, such a Zeitgeber must be linked to the organism’s physiological condition, and it must act as an honest signal, immune to mutations and cheating. This Zeitgeber is likely a finite or limited resource. A putative mechanism may be related to cellular aging, such as a Hayflick limit, which describes the observation that a human cell can only replicate and divide a finite number of times (Chan et al., 2022). However, a Hayflick limit has not been described for insect cell cultures, to the best of our knowledge. Whatever the mechanism, it cannot be causally related to reproduction and caste, and is thus likely rooted in the shared physiology and metabolism.

Kin selection is a powerful force and has overridden individual selection several times, resulting in the major transitions in evolution (Bourke, 2011), including sterile organisms without direct fitness in the holometabolous social insects (Hamilton, 1964). The discovery of continousparous reproduction, coupled with the observation that queens and workers show overlapping patterns in the pace and shape of aging, lead us to hypothesize that kin selection underlies the evolution of programmed aging in *C. obscurior*. Our hypothesis is explicitly distinct from the idea of “phenoptosis” i.e. the removal of old individuals for the betterment of the species. Instead, we propose that programmed aging has evolved to optimize investment into reproduction in a superorganism.

## Glossary

Aging: Time passing by. Neutral in terms of fitness - often confused with senescence.
Life expectancy: Species (mean) lifespan estimate based on mortality rates.
Lifespan: The maximum age an organism can reach - often confused with life expectancy.
Mortality rate (age-specific): Expected probability to die at a given age in a particular population.
Pace of aging: Related to the extension of the life of an organism (either mean or maximum lifespan) See Baudisch, 2011.
Senescence: When an organism shows signs of increasing mortality and decreasing fertility.
Shape of aging: Mortality and fertility of an organism for each time point, standardized by age.

## Acknowledgments

Thanks to B. Dofka, M. Schlossberger, L. Mecka, and M. Fallböhmer for help feeding colonies. This research was supported by the Deutsche Forschungsgemeinschaft with a grant to JO and JH (DFG; OE549/2-2,3), within the frame of the DFG Research Unit FOR2281. No funding sources were involved in study design, data collection and interpretation, or the decision to submit the work for publication.

